# Integrative methylome and transcriptome analysis of Japanese flounder (*Paralichthys olivaceus*) skeletal muscle during development

**DOI:** 10.1101/516187

**Authors:** Jingru Zhang, Shuxian Wu, Yajuan Huang, Haishen Wen, Meizhao Zhang, Jifang Li, Yun Li, Xin Qi, Feng He

**Author notes:** These authors contributed equally to this work (Shuxian Wu, Yajuan Huang, Haishen Wen, Meizhao Zhang, Jifang Li, Yun Li, Xin Qi).

## Abstract

DNA methylation is an important epigenetic modification in vertebrate and is essential for epigenetic gene regulation in skeletal muscle development. We showed the genome-wide DNA methylation profile in skeletal muscle tissue of larval 7dph (JP1), juvenile 90dph (JP2), adult female 24 months (JP3) and adult male 24 months (JP4) Japanese flounder. The distribution and levels of methylated DNA within genomic features (1stexons, gene body, introns, TSS200, TSS1500 and intergenic) show different developmental landscapes. We also successfully identified differentially methylated regions (DMRs) and different methylated genes (DMGs) through a comparative analysis, indicating that DMR in gene body, intron and intergenic regions were more compared to other regions of all DNA elements. A gene ontology analysis indicated that the DMGs were mainly related to regulation of skeletal muscle fiber development process, Axon guidance, Adherens junction, and some ATPase activity. Methylome and transcriptome clearly revealed a exhibit a negative correlation. And integration analysis revealed a total of 425, 398 and 429 negatively correlated genes with methylation in the JP2_VS_JP1, JP3_VS_JP1 and JP4_VS_JP1 comparison groups, respectively. And these genes were functionally associated with pathways including Adherens junction, Axon guidance, Focal adhesion, cell junctions, Actin cytoskeleton and Wnt signaling pathways. In addition, we validated the MethylRAD results by bisulfite sequencing PCR (BSP) in some of the differentially methylated skeletal muscle growth-related genes (Myod1, Six1 and Ctnnb1). In this study, we have generated the genome-wide profile of methylome and transcriptome in Japanese flounder for the first time, and our results bring new insights into the epigenetic regulation of developmental processes in Japanese flounder. This study contributes to the knowledge on epigenetics in vertebrates.

**Author summary:** Epigenetic mechanisms like DNA methylation have recently reported as vital regulators of some species skeletal muscle development through the control of genes related to growth. To date, although genome-wide DNA methylation profiles of many organisms have been reported and the Japanese flounder reference genome and whole transcriptome data are publically available, the methylation pattern of Japanese flounder skeletal muscle tissue remains minimally studied and the global DNA methylation data are yet to be known. Here we investigated the genome-wide DNA methylation patterns in Japanese flounder, throughout its development. These findings help to enrich research in molecular and developmental biology in vertebrates.

## Introduction

As to genetic regulations, a growing number of studies have reported that epigenetic modifications play critical roles in gene expression. Epigenetics refers to heritable changes that modify DNA or associated proteins but without changing the fundamental DNA sequence itself [1]. The epigenome is a dynamic entity influenced by predetermined genetic programs and external environmental cues [2]. DNA methylation is an imporant epigenetic modification of the genome found in most eukaryotes and plays a key role in muscle development. It occurs at the C5 position of cytosine within CpG and non-CpG in the genome. The regulation and mechanisms of the DNA methylation still remain enigmatic, although it is essential for normal development and crucial in many biological processes, such as gene expression regulation,embryogenesis, cellular differentiation, genomic imprinting, X-chromosome inactivation, maintenance of genomic stability by transposon silencing [3–6]. Now, DNA methylation has attracted much attention owing to its broad impact, reversibility, heritability and genetic characteristics.

The profile of DNA methylation across the genome is important to understand DNA methylation dynamics during different developmental muscle development. The genome-wide DNA methylation profiles and functional analysis of many organisms, such as human [7], rat [8], Arabidopsis [9] has been reported. However, little is known about the DNA methylation patterns in Japanese flounder.

Japanese flounder is one of the commercially important marine fish in China and has been widely cultured in recent years. Skeletal muscle represents the most abundant tissue in the body and its features have a direct impact on meat quality. Understanding the growth and development of skeletal muscle is important. Skeletal muscle development is a very complicated but precisely regulated process, which contains four steps: determination of myoblasts, proliferation of myoblasts, differentiation and fusion of myoblasts into myotubes and myofibers, and growth and maturation until postnatal [10–11]. The research of complex mechanism underlying skeletal muscle development is helpful to genetic improvement for meat quality. In Japanese flounder, the muscle mass and meat quality are mostly determined by the size and the number of myofibers. Hence, we chose three postnatal stages (larval, juvenile and adult stage) which are key points in Japanese flounder skeletal muscle growth and development. The comprehensive analyses of these specific stages should help to understand the developmental characteristics in Japanese flounder skeletal muscle.

Many previous studies have concentrated on the impacts of DNA methylation and even located sites in the promoter or the first exon of a gene. It generally leads to transcriptional silencing and suppresses the corresponding protein products in most eukaryotes [6,12–14]. Thus, DNA methylation plays critical roles in cellular processes and the development of skeletal muscle tissue.

There are many approaches to decipher a genome-wide DNA methylation profile, including Methylation-dependent restriction-site associated DNA sequencing (MethyIRAD), MeDIP-seq and whole-genome bisulfite sequencing (WGBS). The gold standard to determine the DNA methylome is genome-wide bisulfite sequencing, which firstly converts all the unmethylated cytosines into uracil while left the methylated cytosines unchanged by sodium bisulfite under denaturing conditions, which can be distinguished subsequently by sequencing[15]. <u>whereas</u> genome-wide bisulfite sequencing is highly expensive and time-consuming. Now, studies have shown that MethylRAD is a suitable method for high-throughput sequencing to analyze the DNA methylation status of methylated genome regions at a fraction of the cost and time of genome-wide bisulfite sequencing. MethylRAD uses methylation-dependent restriction enzymes which can specifically discriminate methylated cytosines between CG and non-CG methylation. These enzymes have the unique ability to produce 32-base-long fragments around fully methylated restriction sites, which are suitable for high-throughput sequencing to profile cytosine methylation on a genomic scale [16,17]. Many recent studies have shown that MethyIRAD can reflect the relative genome-wide DNA methylation profile [18,19]. Therefore, we chose MethyIRAD to analyze genome-wide profiles of DNA methylation in Japanese flounder in our study.

In this study, we have performed the first integrated genome-wide analysis of DNA methylation, and mRNA transcriptional activity, using the transcriptome and MethylRAD (a simple genomic methylation site detection method) [16,20] data by high-throughput, deep-sequencing technologies and subsequent bioinformatics analysis. A series of genes involved in the development of skeletal muscle were confirmed to show simultaneously differential expression levels and DNA methylation levels. These findings provided comprehensive insights into the skeletal muscle development during different developmental stages. Skeletal muscle tissues were used in this study, namely, the larval 7dph (JP1), juvenile about 90dph (JP2), adult female about 24 months (JP3) and adult male bout 24 months (JP4).

## Results

### Global mapping of DNA methylation in Japanese flounder

The MethylRAD analysis was used to study the global mapping of DNA methylation pattern in the Japanese flounder skeletal muscle tissues of JP1, JP2, JP3 and JP4. We generated about 439 - 751 million raw reads from each samples. After low-quality data filtration, about 42 to 54 million reads assessed as clean data were analyzed and mapped (S1 Table). Base distribution and quality distribution maps of clean reads were plotted (S1 Fig). Of the high-quality methylation tag libraries in the twelve samples, 71.76-81.08% were comparable to unique positions with a high-quality read alignment against the Japanese flounder reference genome using SOAP software (version 2.21) (S1 Table).

The percentage of the DNA methylation sites (CCGG sites and CCWGG sites) in each sample are shown (S2 Fig). We found a substantial amount of CCGG methylation and a small amount of CCWGG methylation. Therefore, we analyzed the genome coverage of the CCGG, CCWGG sites under different sequencing depth (S2 Table). The sequencing depths of the DNA methylation sites (CCGG sites and CCWGG sites) in each sample are shown in a box plot (the number of methylation sites in each sample had a depth higher than 3) (Fig 1). The JP1, JP2, JP3 and JP4 methylation sites (CCGG/CCWGG) were identified on the chromosomes of Japanese flounder. Methyl-RAD reads were detected in most chromosomal regions (chromosomes 1–24) in each group (Figure 2).

**Figure 2.**
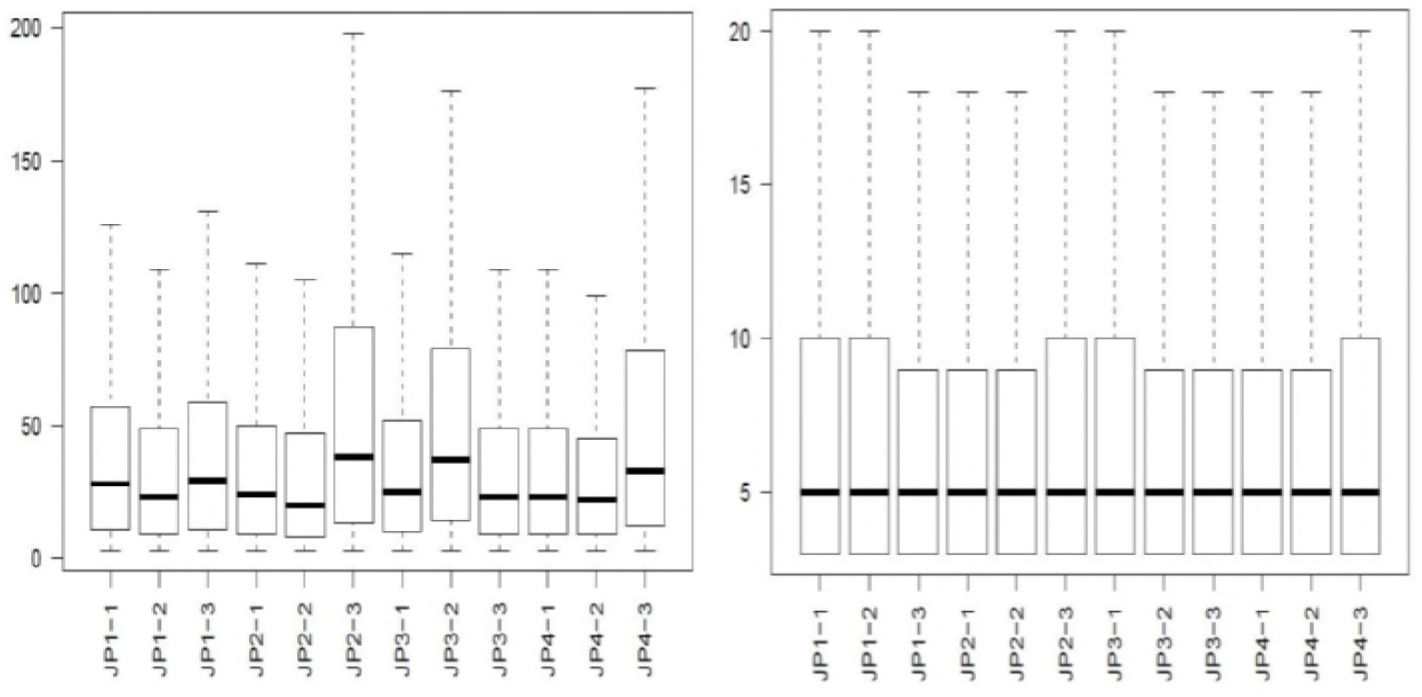
The distribution of DNA methylation sites in the CCGG and CCWGG on chromosomes. The distribution of DNA methylation sites on chromosomes 1 to 24 of the Japanese flounder genome is shown for each sample. From the outside to the inside is JP1, JP2, JP3, JP4 respectively.

### Distribution of DNA methylation sites of different functional regions

The distribution of MethyIRAD reads that were aligned on a unique locus in different genome regions represents a genome-wide methylation pattern. We obtained the DNA methylation site annotation of the Japanese flounder genome and the comparison of average methylation sites showed that there were differential methylation site distribution in different components of the genome. The distribution patterns of most methylated sites at the different elements of genomes were similar in the four groups. We found that the major proportion of DNA methylation sites were mainly enriched in the intergenic regions followed by the regions at the gene body, TSS1500 (upstream 1500 bp of transcription start sites TSS), intron, 1stexon and TSS200 (upstream 200 bp of TSS) at both the CCGG and CCWGG sites (Fig3A). Among all the classes, the average methylation sites of promoter was the lowest. Furthermore, all gene regions in the JP1, JP3 and JP4 group exhibited higher number of methylation site than those in the JP2 group, while JP3 group showed lower number of methylation site than those in the JP4 group (Fig 3B). The distribution of methylation sites on different gene elements in each sample indicates that the skeletal muscle growth difference during different developmental stages might be associated with global methylation.

**Figure 3.**
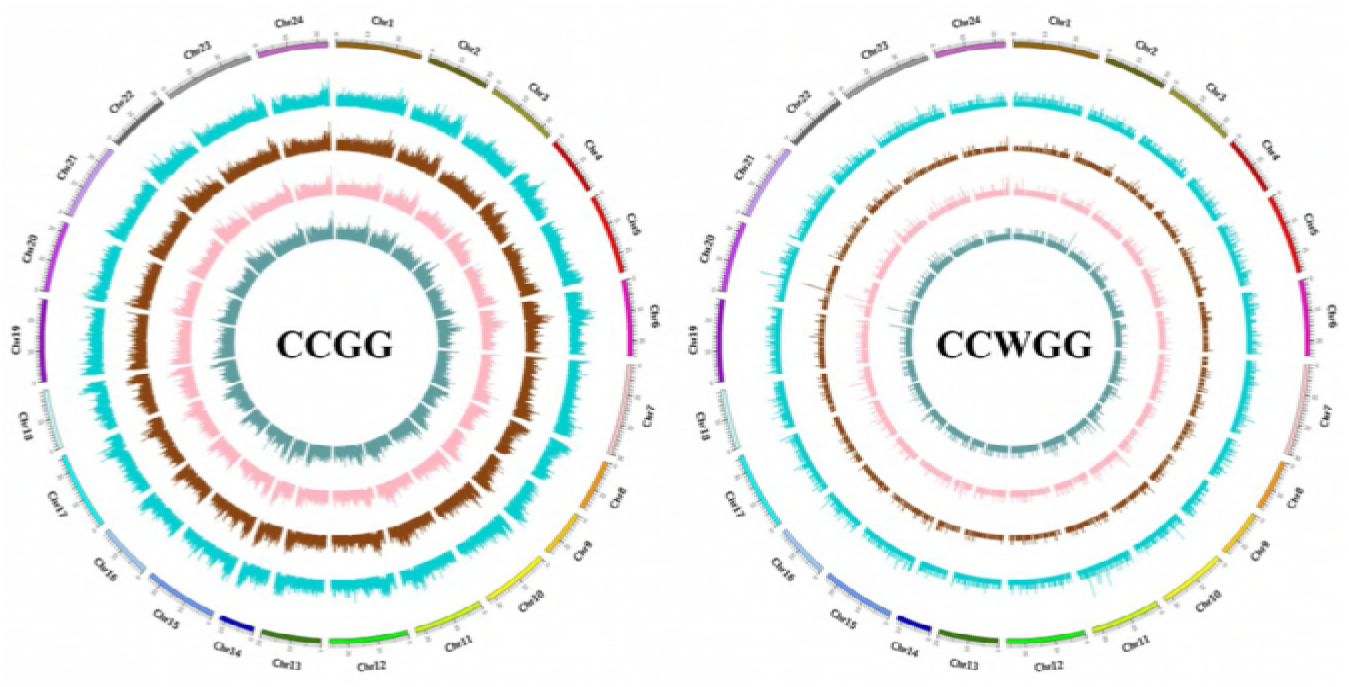
**(A) DNA methylation site on different gene function components distribution histogram**. 1stexon: the regions of the first exon; Body: the whole exons of genes(except 1stexon); TSS200: the upstream 200 bp of the transcription termination site (TSS); TSS1500: the upstream 1500 bp of TSS; intron:the whole introns of genes ;“intergenic” indicated the intergenic regions (CCGG methylation sites is shown, left and CCWGG methylation sites is shown, right). The y-axis shows the number of methylation sites. The x-axis shows the different components of the genome. **(B) DNA methylation distributions of CCGG and CCWGG sites with differential methylation levels**. The y-axis shows the number of methylation sites. The x-axis shows the CCGG, CCWGG sites and the whole CCGG, CCWGG sites.

### Relative quantification of DNA methylation levels around the Gene body

We found that the DNA methylation site distribution curve had TSS representing an upstream sequence centered on the transcription initiation site, and TTS representing a downstream sequence centered on the transcription termination site. Hence, we analyzed the distribution of DNA methylation in the 2 kb region upstream of the TSS, gene body (the entire gene from the TSS to the transcription termination site (TTS). The region around the TSS is crucial for gene expression regulation. The DNA methylation level dramatically decreased in the 2 kb region upstream of the transcription start sites (TSS) and dropped to the lowest point before the TSS and increased sharply towards the gene body regions and stayed at a plateau until the TTS. The DNA methylation levels at either the CCGG sites or the CCWGG sites in the gene regions were similar for four groups (Fig 4).

**Fig 4.**
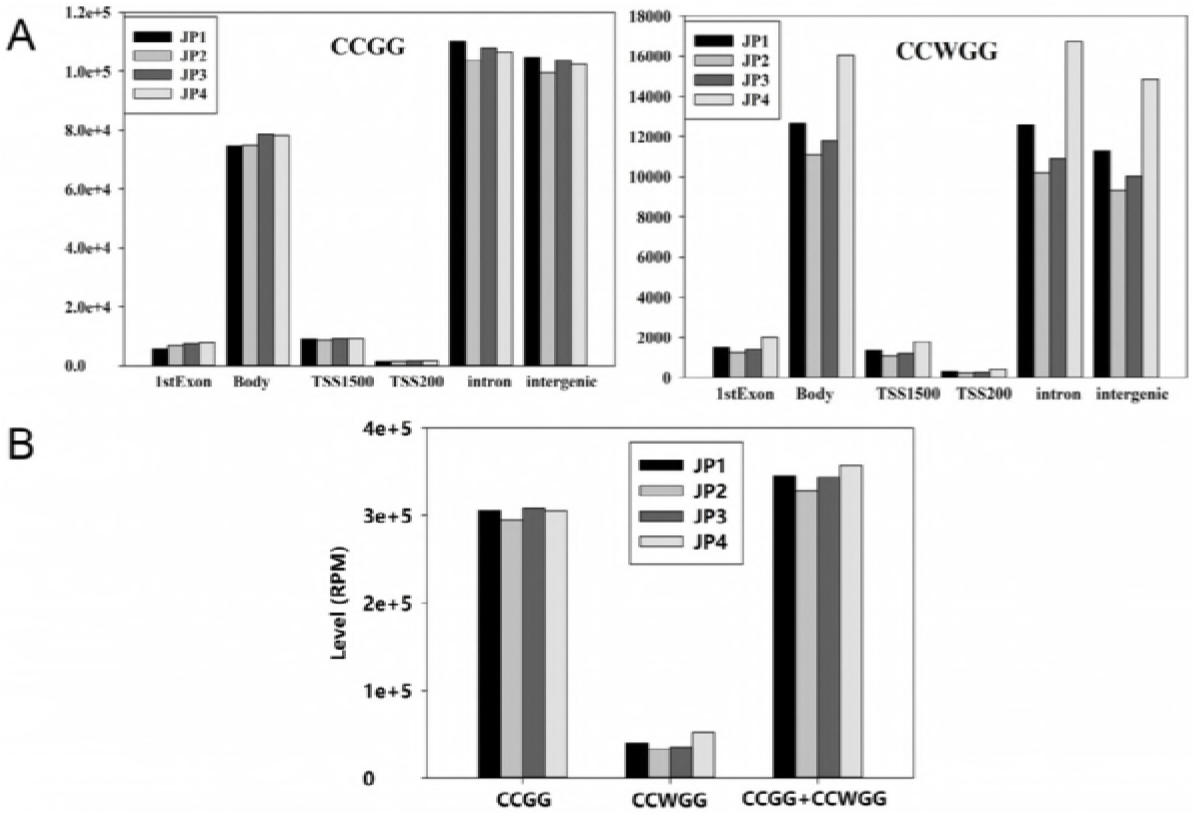
Distribution of Methyl-RAD reads around gene bodies. The x axis indicates the position around gene bodies, and the y axis indicates the normalized read number. This figure reflects the methylation level around gene bodies. CCGG sites methylation level is shown, left and CCWGG sites methylation level is shown, right.

### Differentially methylated regions (DMRs) analysis

To characterize the differences of DNA methylation levels among samples, DMRs were detected. For assessing the methylation level of differential methylation sites between four groups for the three biological replicates, the cluster heat map was shown to further show the changes in CCGG/CCWGG methylation levels among the groups. Hypomethylated CCGG/ CCWGG sites in samples are clustered at the bottom, whereas hypermethylated CCGG/CCWGG in samples are massed on upper cluster heat map. Interestingly, hierarchical cluster analysis results indicated that there were unique methylation patterns among four groups, and showed distinctive interindividual and intraindividual differences in methylation profiles among groups (S3 Fig).

The number of hypermethylation DMRs is less than hypomethylation DMRs. The number of DMRs in CCGG sites is lower than that in CCWGG sites. DMRs that are unique or shared among the four groups are shown (Fig 5). The results of a box-plot analysis of DMRs showed that the methylation level of the JP2 group was the lowest among four groups and the JP3 group is lower than that in JP4 group (S4 Fig). The pie map distribution of differential methylation sites with differential methylation levels on different functional components was drawn according to the positional information of the differentially methylated site-related gene, and the results are shown (S5 Fig). The results showed that the CCGG/CCWGG site were mostly enriched in the intergenic regions, followed by the gene coding regions (1stExon + other extrons), the TSS1500 (upstream 1500 bp of TSS), intron and TSS200 (upstream 200 bp of TSS) (S5 Fig).

**Fig 5.**
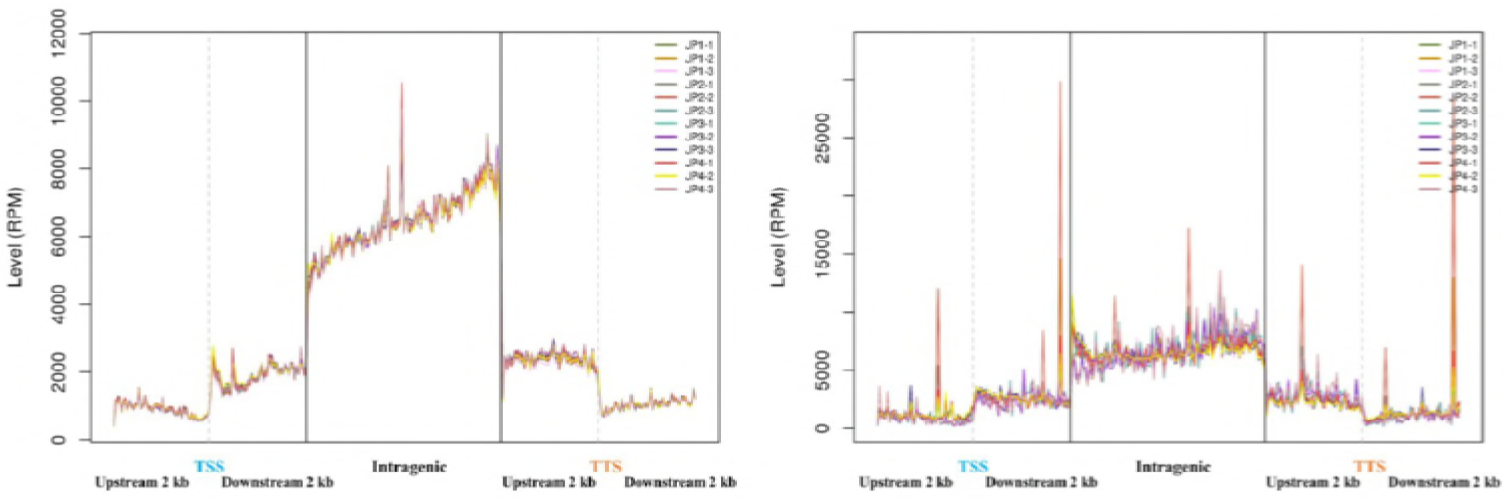
The Venn diagram for comparison of DMRs that are unique or shared in four groups derived from Japanese flounder. For CG context methylation, certain methylated C sites are defined as hypermethylation sites, at which the methylation level is over 75%; and some others are defined as sites of hypomethylation, at which the methylation level is less than 75%. For non-CG context methylation, hyper- and hypomethylation sites are defined as those at which the methylation levels are over or under 25%, respectively.

### Analysis of differential methylation site-related genes

To investigate the differential methylation site-related genes regulatory role, the function of a gene was described by the GO and KEGG enrichment analysis of the gene where the differential DNA methylation sites were located. We found that these methylation site-related genes were significantly enriched in some biological processes and signaling pathway important for skeletal muscle development. The GO enrichment analysis top30 bar graph is shown (S6 Fig). The “Focal adhesion”, “actin cytoskeleton”, “Adherens junction”, “**cell junctions**”, “**Wnt signaling pathways**”, “Axon guidance”, “Wnt signaling pathway” and “Hippo signaling pathway” were significantly enriched in Japanese flounder (S7 Fig).

### MethylRAD-seq data validation by bisulfite sequencing

To validate the results obtained with MethyIRAD-seq data, according to the GO and KEGG enrichment analysis of the DMGs, three genes (Myod1, Six1 and Ctnnb1) related to skeletal muscle developmet were selected from MethyIRAD-Seq data in the Japanese flounder genome for analysis by bisulfite sequencing. Ctnnb1 was up-methylated in the JP2_VS_JP1, JP3_VS_JP1 and JP4_VS_JP1 comparison groups, respectively; Six1 was up-methylated in the JP3_VS_JP1 and JP4_VS_JP1 comparison groups, respectively; and MyoD1 was up-methylated in the JP4_VS_JP3 comparison group and as down-methylated in the JP3_VS_JP2 comparison group. The bisulfite sequencing results showed a high degree of consistency with the MethylRAD data (Fig 6). These results indicated that our genome-wide methylation results obtained by MethylRAD are reliable.

**Fig 6.**
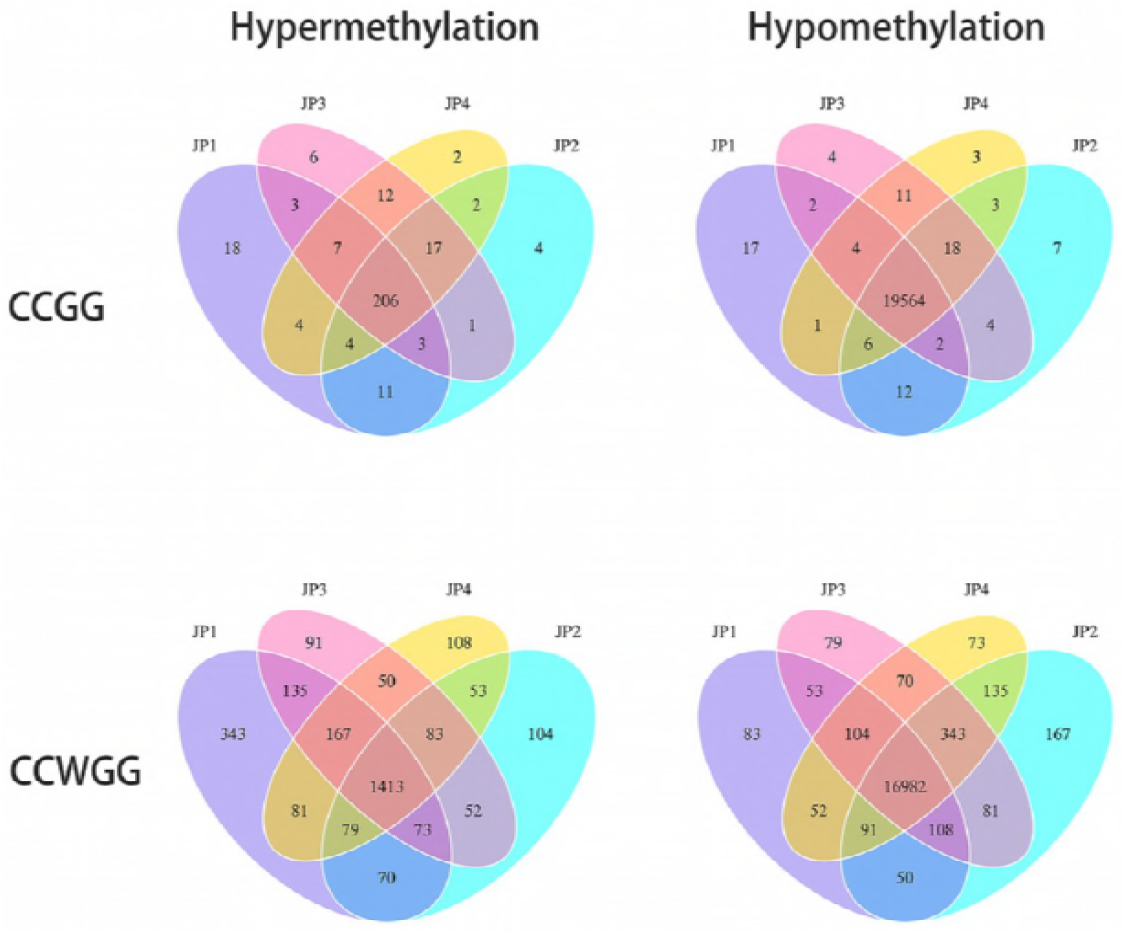
The validation of MethylRAD data by bisulfite sequencing (BSP). Three genes obtained from MethylRAD data was selected randomly and its methylation pattern was profiled by BSP. The box indicated amplification regions. CpG dinucleotides are represented by circles on vertical bars. Each line represents an independent clone, and methylated CpGs are marked by filled circles, unmethylated CpGs by open circles. Average methylation was calculated for all CpG sites in each stage. There were three samples in each group, respectively and for each sample typically 10 clones were used to show DNA methylation levels. Different colors in the right show different methylation level.

### Transcriptome assembly and annotation

Using RNA-Seq, this study compared the transcriptomic landscapes of skeletal muscle from the larval, juvenile and adult (female and male) stages used to construct mRNA libraries. All the samples sequenced on the Illumina HiSeq X Ten platform and 150 bp paired-end reads were generated. The sequencing reads were analyzed using Tophat software by alignment with the Japanese flounder reference genome. Raw reads were processed using the NGS QC Toolkit to reduce the impact of sequencing errors. After filtering low quality reads, reads containing adapter, reads containing ploy-N, among the aligned reads, a total of 91,134,425 (JP1), 91,104,445 (JP2), 91,857,072 (JP3) and 92,720,615(JP4) average clean reads were mapped, and on average approximately 77.63%(JP1), 71.42%(JP2), 78.50% (JP3) and 76.18% (JP4) of the reads individually were totally mapped to the Japanese flounder genome and approximately 74.67%(JP1), 65.34%(JP2), 65.80%(JP3) and 66.10%(JP4) of the reads in each sample were uniquely mapped to the Japanese flounder genome in each sample. Multiply mapped(JP1:2.96%, JP2:6.09%, JP3:12.69%, JP4:10.08%) reads were excluded from further analyses and other Parameters are presented (S3 Table).

### Comparative and enrichment analysis of differentially expressed genes

We defined genes with fold changes > 2 and P-values < 0.05 were recognized as significantly differentially expressed. Different expression genes that are unique or shared among the four groups examined are shown(Fig7).

**Fig 7.**
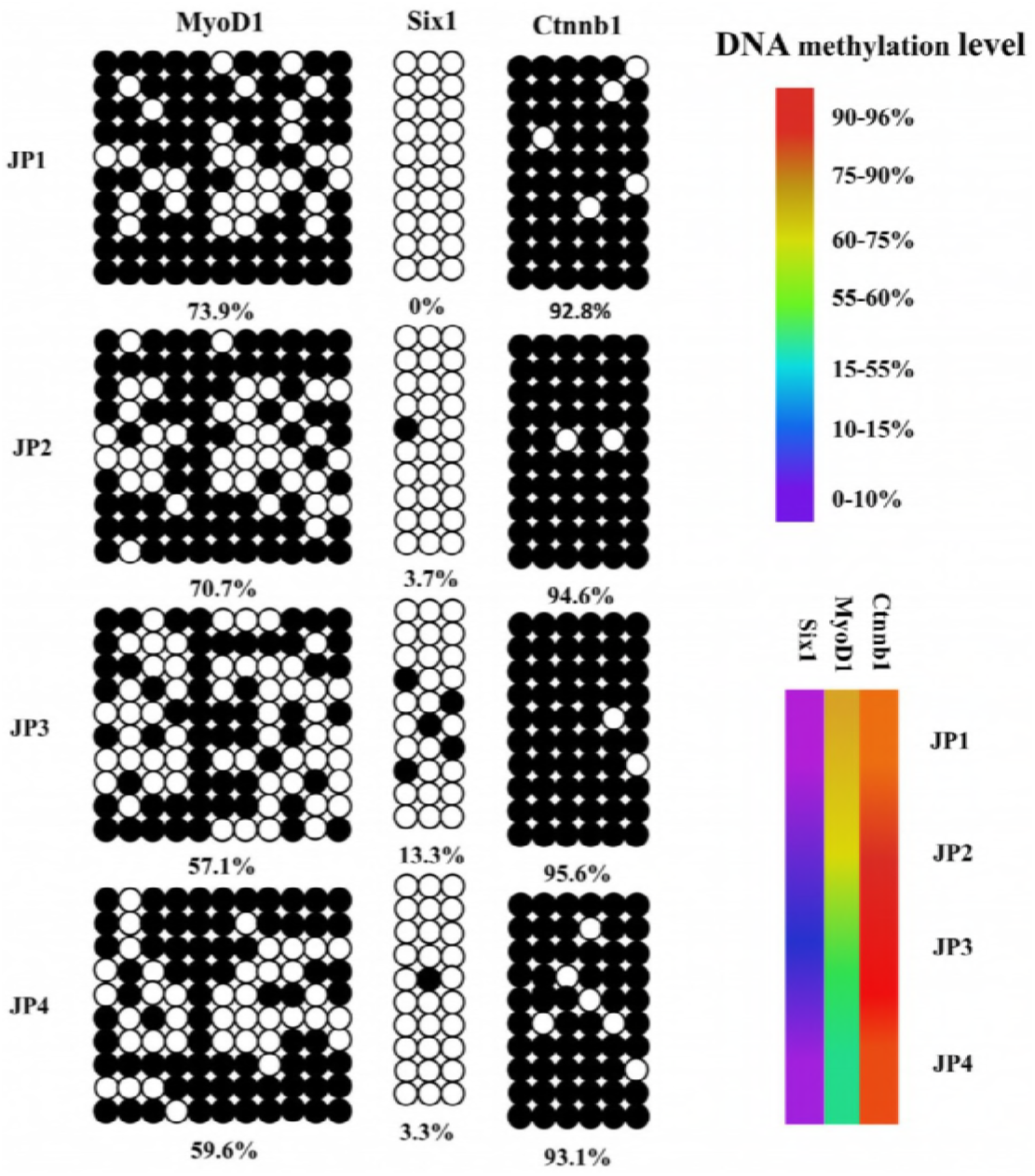
The Venn diagram for comparison of DEGs that are unique or shared in four groups. DEG: Differently methylated genes

The DEGs among groups were conserved and were mainly enriched in the cellular component, molecular function, and biological process categories (S8 Fig). And the analysis of KEGG pathway revealed that multiple pathways involved in growth and development were clearly enriched in the Japanese flounder DEGs, including the “Axon guidance”, “Adherens junction” and “Focal adhesion” also exhibited over-represented in the GO terms targeted by the DMGs (S9 Fig).

### RNA-Seq data validation

To examine the reliability of the RNA-seq results, three DEGs (MyoD1, Six1 and Ctnnb1) involved in the development of skeletal muscle were selected for validation using qRT-PCR. The mRNA expression levels of these key genes, such as Ctnnb1 related to cytoskeleton and cell adhesion, was down-regulated in the JP2_VS_JP1 comparison groups; Six1 corrected with regulation of skeletal muscle cell proliferation and skeletal muscle fiber development and MyoD1 in connection with positive regulation of myobalst differentiation and skeletal muscle cell differentiation, were all up-regulated in the JP2_VS_JP1 comparison groups, respectively. As shown in Fig 8, the qRT-PCR expression patterns of the three DEGs were in agreement with the RNA-seq data.

**Fig 8.**
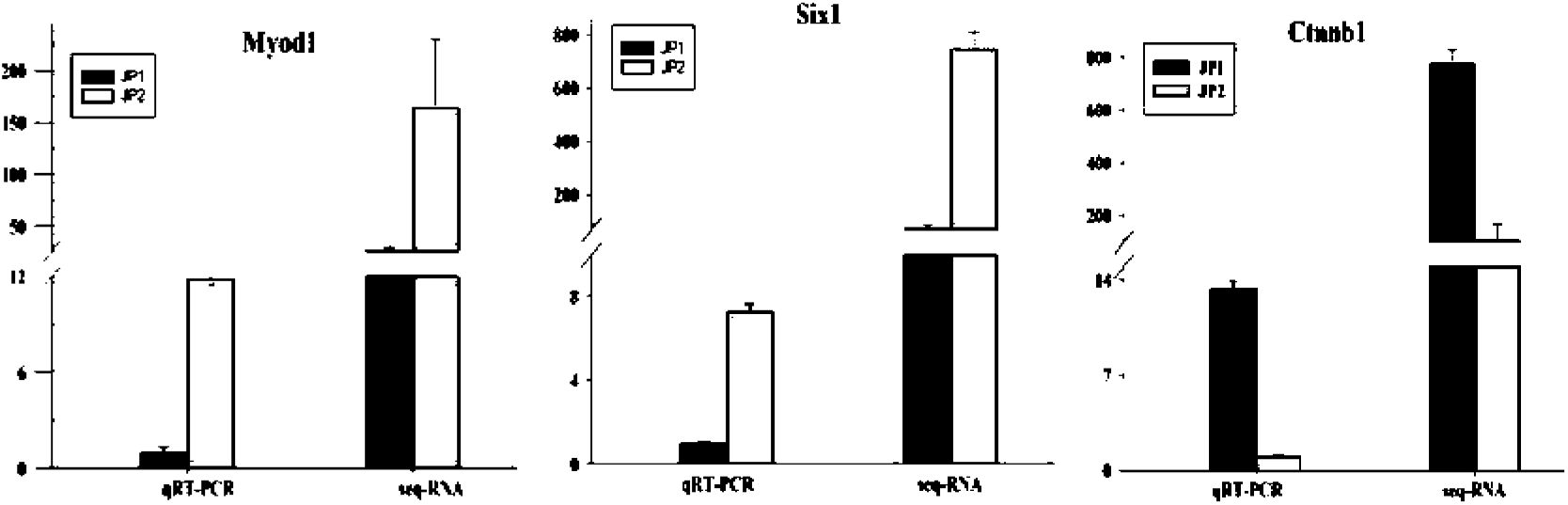
The validation of RNA-seq data by qRT-PCR. Three genes obtained from RNA-seq data was selected and was validated by qRT-PCR. X axis indicated the relative RNA-seq and qRT-PCR results of genes. Y axis indicated the relative expression levels of genes.

### Association analysis of methylRAD and the transcriptome (RNA-Seq)

The association analyses between the transcriptome and methylation were based on RNA-Seq and MethylRAD sequencing data. We calculated each gene’s methylation level and expression level in larval, juvenile and adult female and adult male period Japanese flounder in terms of DNA methylation and the mRNA transcriptome. We observed that methylation levels correlates negatively with expression levels in both the CCGG and CCWGG pattern in larval, juvenile and adult Japanese flounder skeletal muscle tissue (Fig 9B). Furthermore, to explore the relationship between these DMGs and the DEGs found at the transcriptome level, an association analysis was performed. We found that a lot of genes that both different methylated and didfferent expressed in the JP2_VS_JP1, JP3_VS_JP1 and JP4_VS_JP1 comparison groups, respectively. However, only few genes simultaneously showed differential expression levels and DNA methylation levels in the JP2_VS_JP3, JP2_VS_JP4 and JP3_VS_JP4 comparison groups, respectively (Fig 9A and S5 Table). We speculate that DNA methylation mainly affects skeletal muscle development in the larval period, and differences in skeletal muscle between juvenile and adult period Japanese flounder may be affected by other factors rather than DNA methylation, and that require further research.

**Fig 9.**
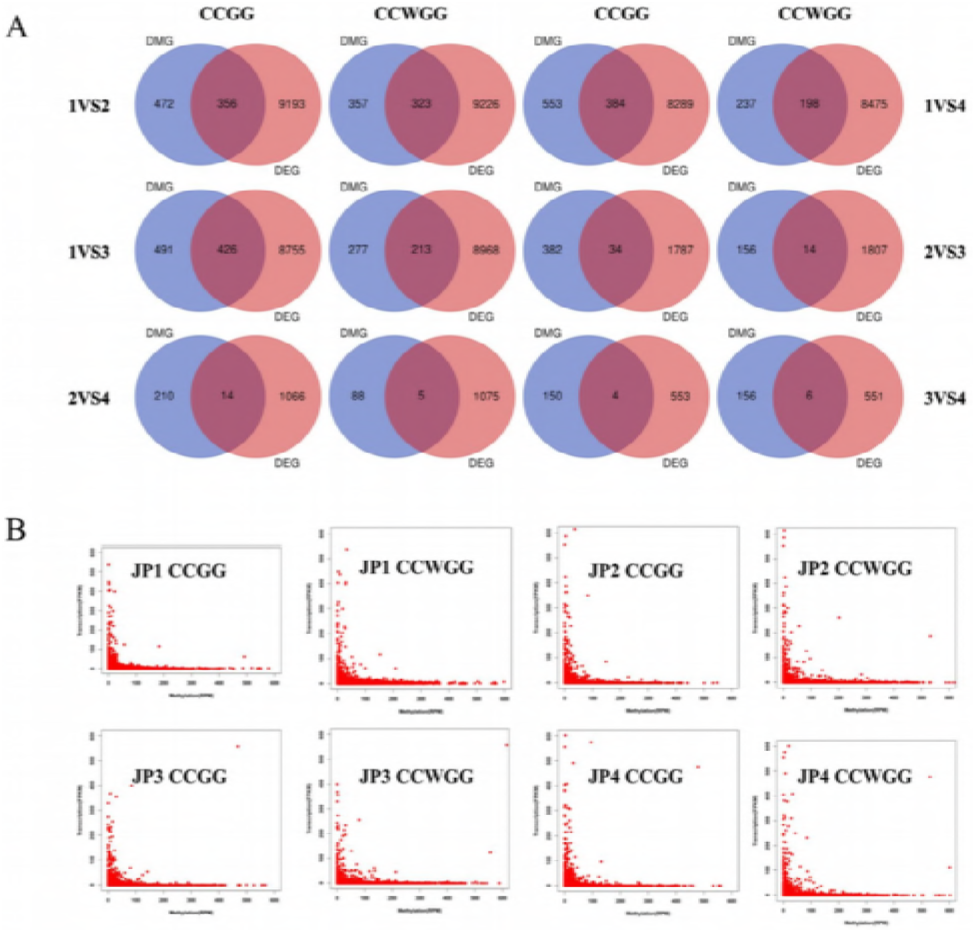
Relationship of differential expression levels of genes and their DNA methylation levels. a indicated the Venn diagrams of genes showing differential expression levels and/or differential DNA methylation levels by pairwise comparison analysis. DMG, differentially methylated gene; DEG, differentially expressed gene. b indicated the distribution characteristics of gene methylation and expression levels. JP1 (larval Japanese flounder); JP2 (juvenile Japanese flounder); JP3 (adult female Japanese flounder); JP4 (adult male Japanese flounder). X axis indicated the relative DNA methylation levels of genes. Y axis indicated the relative expression levels of genes.

Among these DEGs which also exhibited differential methylation levels, a large proportion of them appeared to be negatively correlated with their DNA methylation levels in the JP2_VS_JP1, JP3_VS_JP1 and JP4_VS_JP1 comparison groups, respectively. The results show a total of 238 and 187 negatively correlated genes with methylation in the JP1 and JP2 libraries, a total of 273 and 125 negatively correlated genes with methylation in the JP1 and JP3 libraries, a total of 310 and 119 negatively correlated genes with methylation in the JP1 and JP4 libraries in the CCGG and CCWGG site, respectively. These results suggest that DNA methylation makes a difference in the skeletal muscle during the developmental stages from larval to juvenile and adult Japanese flounder.

In the result, 118 were methylation down-regulated and expression up-regulated and 307 were methylation up-regulated and expression down-regulated in the skeletal muscle during the JP2_VS_JP1 comparison group. 118 were methylation down-regulated and expression up-regulated and 280 were methylation up-regulated and expression down-regulated in the skeletal muscle during the JP3_VS_JP1 comparison group. 132 were methylation down-regulated and expression up-regulated; and 307 were methylation up-regulated and expression down-regulated in the skeletal muscle during the JP4_VS_JP1 comparison group (S5 Table).

To further investigate the signaling pathway associated with negatively correlated genes with methylation and expression levels in juvenile and adult compare to larval period, we performed KEGG enrichment analysis of these genes. The results showed that there were significantly enriched KEGG signaling pathway (P <0.05) between larval and juvenile, between larval and female male adult, and between larval and male adult Japanese flounder, respectively (S10 Fig). In our study, KEGG enrichment analysis screens criteria for pathway entries with the number of differential genes greater than 2, however, there are too few differential genes to show in adult female and adult male compared to juvenile Japanese flounder and between adult female and adult male Japanese flounder, respectively. The pathway terms showing the highest level of significance were the Adherens junction, Axon guidance, Focal adhesion, cell junctions, actin cytoskeleton, Wnt signaling pathways and Hippo signaling pathway involved in the regulation of growth and development of skeletal muscle are shown.

## Discussion

DNA methylation and mRNAs have been studied extensively in the past decades. However, a few studies have focused on Japanese flounder, one of the important economic Mariculture animals. This study is the first to compare systematically the genome-wide skeletal muscle DNA methylation profiles and their relationships to mRNA of larval, juvenile, adult female and adult male Japanese flounder. This study provided a comparative analysis of DNA methylation profiles of Japanese flounder muscle by MethyIRAD. Our data showed almost the entire genome with enough depth to identify differentially methylated regions wite high accuracy and proved that MethyIRAD is a cost-effective approach for comprehensive analyses of the vertebrate genome-wide DNA methylation.

Previous studies shows that DNA methylation is unevenly distributed in genomes, DNA methylation is enriched in the gene body regions, and depleted in the TSS and TTS [21–23]. And the intergenic regions is usually hypermethylated, while promoter regions of genes are relatively hypomethylated compared with the intragenic regions [8,22,24]. Japanese flounder displays analogous methylation pattern with that species. In our analysis, hypermethylation occurred not only at intergentic, gene body regions but also at introns, whereas the promoter (around TSSs) remains hypomethylated. The intracellular hypermethylation in the Japanese flounder genome further indicates that this methylation pattern may be a more conservative mechanism among species. In contrast to previous research in animals [8,25–26], we did not observe a higher methylation level in exons than in introns in Japanese flounder.

DNA methylation is one of the main epigenetic modification mechanisms, the analysis of DMRs within individuals is important. In several studies, different levels of DNA methylation could regulate stage-specific transcription and may be important during development and differentiation [27]. Thus, the analysis of DMRs among stages is essential in understanding stage-specific gene expression. We also observed that distribution of DNA methylation in the four groups showed generally conserved pattern, some DMRs were detected a high density in the intergentic, gene body and introns and a low density in the TSS200, TSS1500 and first exon regions. The first exon contained relatively few DMRs within the gene body, which may be the result of certain motifs overlapping between the promoter and the first exon. DNA methylation, especially intronic DNA methylation, may be associated with alternative splicing [28].

The DNA methylation status of promoter and gene body regions play an important role in the regulation of gene expression regulation via alteration in chromatin structure or transcription elongation efficiency [29–31]. Most of the promoter regions were hypomethylated in the vertebrate genome and one long established role of DNA methylation in gene promoter regions is the repression of gene expression [32,33]. Previous studies have demonstrated that DNA methylation in gene body regions impeded transcription elongation in Human, chicken, Neurospora crassa and Arabidopsis thaliana [22–23,34–35]. Methylation of these elements is known to be a crucial factor in the maintenance of genomic stability through the suppression of transcription, transposition, and recombination [8]. Thus, these results suggest that methylation has important effects on gene transcription in individual with different developmental stages. However, DNA methylation is only one of the regulators that influence gene expression. Since the interactions between transcription factors and methylated DNA could impact gene expression regulation and chromatin remodelling, changes in methylation may affect the expression of a gene [36]. Further studies are needed to explore the complicated epigenetic mechanism underlying growing. In summary, differences in DNA methylation patterns and the status of DMRs in the four groups of different developmental stages may play a crucial role in the process of development and the corresponding gene expression.

In this study, we constructed RNA-seq and DNA methylation libraries from skeletal muscle tissues of different developmental stages using transcriptome sequencing and the MethylRAD methods and discovered some genes simultaneously showed differential expression levels and DNA methylation levels. In our study, we have identified a large proportion of negatively correlated genes in skeletal muscle from larval, juvenile and adult Japanese flounder using deep sequencing technologies. The results showed that there were more methylation up-regulated and expression down-regulated genes increased in the skeletal muscle in the JP2_VS_JP1, JP3_VS_JP1 and JP4_VS_JP1 comparison groups, respectively. A GO enrichment analysis of these negatively corrected genes revealed that the variation in skeletal muscle development was related to biological processes, such as positive regulation of skeletal muscle tissue growth, skeletal muscle fiber development, Adherens junction, cell junctions and axon guidance. Meanwhile, in the present study, the functional annotation indicated that a large proportion of genes were involved in several important signaling pathways, including Wnt signaling pathways, cell adhesion, tight junctions, Adherens junction, hippopotamus signaling pathway, axon guidance, Focal adhesion and cytoskeleton. Axon guidance refers to the process by which growing neural axons follow specific, predictable paths to reach their target locations [37].

Differential methylation changes in this pathway were used as a focus to identify how epigenetic changes during aging could potentially associate with the well-known decrease of skeletal muscle function with increasing age [38]. In addition, we also found the signaling networks that guide diverse cell behaviours and functions are connected to tight junctions transmitting information to and from the cytoskeleton [39], enriched. Previous research showed that the tight junction participated in the regulation of cell growth and differentiation, while adherens junctions participate in contact inhibition of cell growth [40,41]. Several studies, within the last decades, showed that Wnt signaling pathways are involved in myogenesis and regulate muscle formation. In myogenesis, the effect of Wnt signaling leads to the progression of the differentation at early developmental stages and inhibition of this signaling leads to a poor skeletal muscle formation [42–44]. Remodeling of the actin cytoskeleton is critical for mediating changes in many fundamental processes including the cell shape, migration, and adhesion. The regulation of actin cytoskeleton is regulated by a large group of actin binding proteins that modulate actin assembly, disassembly, branching, and bundling, which form actin filament architecture and make it performing various specialized functions [45]. These results have provided direct evidence suggest that DNA methylation may be related to the skeletal muscle development in Japanese flounder. We believe that the differentially methylation of these genes might partially contribute to the Japanese flounder growth difference during different developmental period. However, the epigenetic effects of these genes on Japanese flounder growth still require further study in the future. This study expands the Japanese flounder methylated genes and could initiate further study in the muscle development of Japanese flounder.

In addition, We discovered some differentially DNA methylated genes involved in skeletal muscle development in larval juvenile and adult Japanese flounder. For example, we found that MyoD1, a master regulatory gene of skeletal muscle differentiation [46]. Another well-known gene named myf6 (MRF4) is involved in inducing fibroblasts to differentiate into myoblasts and affects skeletal muscle development [47]. Six1 has been shown to play a pivotal role in skeletal muscle development [48–50] which is a transcription factor essential for embryonic myogenesis and also regulates MyoD1 expression in muscle progenitor cells. In addition, a recent study shows that Six1 contributes to the regeneration of adult muscle by enhancing and maintaining MyoD1 expression in adult muscle satellite cells in addition to its role in embryonic muscle formation [51]. MyoD1 is able to promote the transformation of multipotent stem cells to skeletal muscle by binding and activating the expression of a subset of pre-myogenic mesoderm genes, including Six1[52]. And gene Ctnnb1 modulates skeletal muscle development by acting on transcription factors controlling myogenesis such as MyoD[53].

We believed that the methylation of these genes might partially contribute to the Japanese flounder growth difference. However, the epigenomic regulation of molecular basis of skeletal muscle among different stages of Japanese flounder growth, which contributes to muscle growth-related genes, is still unclear and require further study in the future.

## Conclusions

We have generated the genome-wide profile of DNA methylation in Japanese flounder for the first time, and our results can be used for depth analyses of the roles played by DNA methylation in Japanese flounder and make that enriches research in molecular and developmental biology in vertebrates. Together, the work performed in this study probably aid in searching for epigenetic biomarkers for muscle growth regulation and promoting further development of Japanese flounder as a model organism for muscle research in other vertebrates.

## Materials and methods

### Ethics statement

All experimental procedures and sample collection were conducted according to the guidelines and were approved and supervised by the respective Animal Research and Ethics Committees of Ocean University of China. The field studies did not involve endangered or protected species. The fish were all euthanized by tricaine methanesulfonate (MS-222).

### Experimental fish and data collection

The experimental animals were collected from Donggang District Institute of marine treasures in Rizhao of Shandong province, and were temporary reared in a 500L bucket in seawater in Ocean University of China within the same environment. About 1000 individuals of larval 7dph (about 50 individuals as one sample) (stage JP1), 40 individuals of juvenile about 90dph (stage JP2), and 80 individuals of adult about 24 months (stage JP3 and JP4) were collected. During our experiment and data analysis, the fish of stage JP1 were too small, so we used about 50 individuals as one sample, other groups included three individuals, which were regarded as biological replicates. All fish were sacrificed in compliance with the international guidelines for experimental animals using tricaine methanesulfonate (MS-222). All fresh skeletal muscle samples were collected (In stage A, we cut off redundant tissue and only retain muscle tissue under the microscope) and the tissues were immediately frozen in liquid nitrogen and then stored at −80°C until DNA and RNA extractions.

### DNA sample isolation and MethylRAD library construction and high-throughput sequencing

Genomic DNA of the four Japanese flounder groups was extracted from skeletal muscle tissues with TIANamp Marine Animals DNA Kit (Cat No.DP324-03) according to the manufacturer’s protocol. The MethylRAD tag libraries were constructed in 12 individuals with four groups following the protocol from Wang et al. [16, 54]. The MethylRAD library was prepared by digesting 200 ng genomic DNA for each sample using 4 U of the enzyme FspEI (NEB, USA) at 37 °C for 4 h. Run 4 μl of digested DNA (~50 ng) on 1% garose gel to verify the effectiveness of digestion. FspEI can recognize 5-methylcytosine (5-mC) and 5-hydroxymethylcytosine (5-hmC) in the CmCGG and mCCWGG sites, and generate a double-stranded DNA break on the 30 side of the modified cytosine at a fixed distance (N12/N16).Accordingly, symmetrical DNA methylated sites were bidirectionally cleaved by FspEI to generate 32-base-long fragments. Then, two adaptors were added to the digested DNAs by T4 DNA ligase (NEB, USA), and the ligation products were amplified in 20 μl reactions by specific primers. PCR products were purified using a MinElute PCR Purification Kit (Qiagen) and pooled for sequencing using the Illumina X-ten PE 150 sequencing platform[16]. Base quality values were calculated using a Phred quality score (Q sanger=−10log10p). Input sequencing data before operation and computing were called raw reads. Raw reads were first subject to quality filtering and adaptor trimming. After operation, the data, including adapter reads and low-quality sequences, were removed from raw reads as clean reads.

### DNA methylation data analysis

To improve the accuracy in the following analysis,filtering pair-end sequencing paired clean reads according to the following terms: (i) remove low quality reads (more than 20% of base mass lower than 20), (ii) remove reads containing adapter and (iii) remove sequences containing too many N bases. The clean reads that did not contain the expected FspEI restriction site were further excluded, and the reads containing the methylated CCGG or CCWGG sites, named MethylRAD-tags, were identified. The MethylRAD-tags were subsequently aligned against the reference genome of Japanese flounder (http://ftp.ncbi.nlm.nih.gov/genomes/all/GCF/001/970/005/GCF_001970005.1_Flounder_ref_guided_V1.0/GCF_001970005.1_Flounder_ref_guided_V1.0_genomic.fna.gz) by SOAP program (version 2.21, parameters: -M4-v2-r0) [55] with two mismatches allowed.DNA methylation sites with a sequence depth of no less than 3 were judged to be reliable.

The distribution and density of methylated cytosine sites on chromosomes were calculated. Furthermore, the distributions of the methylated cytosine sites on different elements of the gene region were evaluated. For relative quantification of MethylRAD data, the DNA methylation levels of the genes were then evaluated by summing the methylation levels of sites that were localized in the gene regions and were determined using the normalized read depth (reads per million, RPM) for each site. (RPM = (read coverage per site/high-quality reads per library) × 1,000,000). The correlation between samples of methylation levels was assessed using Pearson’s correlation coefficient. Upstream and downstream of 2kb sections of the gene body, TSS and TTS were selected and summarized the DNA methylation level of the distribution trend of sequencing reads. In our study, for CG context methylation, certain methylated C sites are defined as hypermethylation sites, at which the methylation level is over 75%; and some others are defined as sites of hypomethylation, at which the methylation level is less than 75%. For non-CG context methylation, hyper- and hypomethylation sites are defined as those at which the methylation levels are over or under 25%, respectively.

The change in methylation level was assessed based on the sequencing depth information of each site in the relative quantitative results of methylation, using R package edge R [56]. A p-value <0.05 and log2FC >1 were considered statistically significant. The function of the gene was described by a GO and KEGG function enrichment analysis of the gene where the differential methylation site was located. The number of genes included in each GO entry and KEGG pathway was counted and the significance of gene enrichment for each GO entry and KEGG pathway was calculated using the hypergeometric distribution test [57], GO entries with the number of corresponding genes greater than 2 in three categories were screened and the GO enrichment analysis results. Differences were considered significant at P < 0.05.

### RNA library construction and high-throughput sequencing

Total RNAs were extracted using TRIzol reagent (Invitrogen, CA, USA) according to the manufacturer’s protocol from the same Japanese flounder as in MethyIRAD analysis. RNA purity and quantification were evaluated using a NanoDrop ND-2000 spectrophotometer (Thermo Scientific). RNA integrity was assessed using the Agilent 2100 Bioanalyzer (Agilent Technologies, Santa Clara, CA, USA). RNAs with high purity were used in library construction by TruSeq Stranded Total RNA with Ribo-Zero Gold (illumina, Cat.No. RS-122-2301) according to the manufacturer’s instructions. These libraries were used for sequencing analysis with Illumina HiSeq X Ten platform and 150 bp paired-end reads were generated. To ensure the reliability of the sequencing data, three fish were used to construct sequencing library during each developmental stage. In total, twelve RNA libraries were constructed and then sequenced with three technological replicates.

### Transcriptome data analysis and functional annotation

Clean reads were generated by filtering the low-quality reads, reads containing adapter, reads containing ploy-N from the raw reads of fastq form by NGS QC Toolkit [58]. All clean reads with high quality were annotated and classified by mapping them to the Japanese flounder reference genome by Tophat (http://tophat.cbcb.umd.edu/). The expressed genes were confirmed based on the annotation information of the clean reads. The expression level of each gene was calculated and normalized by the fragments per kilobase of transcript sequence per million base pairs sequenced (FPKM) [59] using bowtie2 [60] and eXpress (v1.5.1) software [61].

Differential expression analysis of the genes was performed by using the DESeq R package (2012). The NB (negative binomial distribution test) was used to test the difference in the number of reads. The transcript expression was estimated by the basemean value. The significantly DEGs between the two arbitrary samples were identified based on the following thresholds: fold changes > 2 and P-values < 0.05. The assembled transcripts were annotated by Genomes (KEGG) and Gene Ontology (GO). The relevant biological process, cellular component and molecular function of the GO categories and KEGG biological pathways were identified through gene enrichment analyses [62]. The hypergeometric test was conducted to identify the significantly enriched GO terms and KEGG pathway (corrected p-value < 0.05).

### Quantitative RT-PCR

The differential expression patterns of the genes detected by transcriptome data were validated by qRT-PCR analysis. The specific primer pairs were designed for the detection of corresponding genes (S6 Table). The 18S gene from Japanese flounder was selected as internal control. TB GreenTM Premix Ex TaqTM II (TliRNaseH Plus) (Takara, Japan, Codeno. RR820A) in the StepOnePlus Real-Time PCR System was used in the experiments. The relative expression levels of the genes were calculated by the comparative 2-ΔΔCT method according to the manufacturer’s recommendations [63]. Three sample from each developmental stage were used, and three technological replicates were performed to ensure the reliability of quantitative analysis.

### MethylRAD data validation via BSP

To validate the results obtained with MethylRAD data, three different methylated genes among developmental stages reletated to skeletal muscle growth were selected in the Japanese flounder genome for analysis by bisulfite sequencing. Three pairs of bisulfite sequencing PCR (BSP) Primers were designed with Oligo 6.0 (http://www.urogene.org/cgi-bin/methprimer/methprimer.cgi) according to the known sequences (S7 Table). Genomic DNA was extracted from muscle samples at different developmental stages using Marine Animal DNA Kit (TransGen, Beijing, China) following the manufacturer’s instructions. The concentration and purity of DNA were measured by the nucleic acid analyzer Biodropsis BD-1000 (OSTC, China), and the integrity of DNA was evaluated by agarose gel electrophoresis. The Genomic DNA was stored at -20°C for later use. In each developmental stage three fish were used to process the bisulfite modification. Bisulfite modification of 200 ng of genomic DNA was performed using the BisulFlash DNA Modification Kit (EpiGentek, USA) by standard methods. The bisulfite-treated DNA was amplified by PCR with BSP specific primer pair. After a hot start, PCRs were carried out for 40 cycles of 94°C for 40 sec, 50-55°C for 40 sec, and 72°C for 40 sec. PCR products were separated on a 1.5% agarose gel, purified with the TIANGEN gel extraction kit and cloned into the pEASY-T1 vector (TransGen, Beijing, China) and transferred into Trans1-T1 Phage Resistant Chemically Competent Cell (TransGen, China). About ten typically positive clones were selected for each gene and subsequently sequenced to determine the methylation level.

## Supporting information

**S1 Fig. Base distribution and quality distribution maps of clean reads**.

**(PDF)**

**S2 Fig. Comparison of DNA methylation patterns in the four groups**.

**(PDF)**

**S3 Fig. Hierarchical cluster analysis heat-map of differential methylation sites between groups**.

**(PDF)**

**S4 Fig. Methylation levels of DMRs in different groups**.

**(PDF)**

**S5 Fig. The distribution of DMR regions of different functional components**.

**(PDF)**

**S6 Fig. GO enrichment analysis of differential methylation site-related genes on top30 bar graph**.

**(PDF)**

**S7 Fig. KEGG enrichment analysis of differential methylation site-related genes on top20 Signaling pathway**.

**(PDF)**

**S8 Fig. Gene ontology classification of differentially expressed unigenes among groups**.

**(PDF)**

**S9 Fig. KEGG enrichment top20 bubble chart among groups**.

**(PDF)**

**S10 Fig. Annotations and The functional enrichment of genes with significantly negative correlated methylation and expression levels**.

**(PDF)**

**S1 Table. Sample sequencing data volume and match rate**.

**(PDF)**

**S2 Table. DNA methylation site coverage depth in each sample**.

**(PDF)**

**S3 Table. The statistics of Reference genome comparison rate**.

**(PDF)**

**S4 Table. The analysis of differentially expressed genes among the four groups**.

**(PDF)**

**S5 Table. Statistics of differentially methylated and expressed genes**.

**(PDF)**

**S6 Table. Nucleotide sequences of primers used for Real Time PCR in the experiment**.

**(PDF)**

**S7 Table. Primers used for bisulphate PCR(BS-PCR)**.

**(PDF)**

## Acknowledgments

We thank editors and reviewers for comments on the manuscript.

## Author Contributions

**Conceptualization**: Jingru Zhang, Feng He.

**Funding acquisition**: Feng He.

**Investigation**: Shuxian Wu, Yajuan Huang, Haishen Wen, Meizhao Zhang,Jifang Li,Yun Li,Xin Qi.

**Writing – original draft**: Jingru Zhang.

**Writing – review & editing**: Jingru Zhang, Feng He.

---

**Fig 1.** Sequencing depth box diagram of methylation site (CCGG and CCWGG) in each sample. In the box diagram, the box part is the main body of the box-shaped chart, and the middle of the black horizontal line is the median of the data; the upper and lower sides of the box is a quarter of the data that are greater than the upper quartile (Q3), and a quarter of the data are less than the lower quartile (Q1). The interval between Q1 and Q3 is called the inter-quartile range (IQR). The longitudinal lines in the upper and lower sides of the box are tentacle lines. The upper cut off line of the tentacle line is “Q3+1.5 * IQR” and the lower one is “Q11.5 * IQR”.

## References

1. Egger G, Liang G, Aparicio A, Jones PA. Epigenetics in human disease and prospects for epigenetic therapy. Nature. 2004;429(6990):457–63.

2. Thomas A. Down, Vardhman K. Rakyan, Daniel J. Turner, Paul Flicek, Heng Li, Eugene Kulesha et al. A Bayesian deconvolution strategy for immunoprecipitation-based DNA methylome analysis. Nat Biotechnol. 2008 Jul: 26(7): 779–785.

3. Sasaki H, Allen ND, Surani MA: DNA methylation and genomic imprinting in mammals. EXS. 1993; 64:469–486.

4. Courtier B, Heard E, Avner P: Xce haplotypes show modified methylation in a region of the active X chromosome lying 3‗ to Xist. Proc Natl Acad Sci U S A 1995,92(8):3531–3535.

5. Siegfried Z, Eden S, Mendelsohn M, Feng X, Tsuberi BZ, Cedar H: DNA methylation represses transcription in vivo. Nat Genet. 1999;22(2):203–206.

6. Bird A: DNA methylation patterns and epigenetic memory. Genes Dev. 2002; 16(1):6–21.

7. Eckhardt F, Lewin J, Cortese R, Rakyan VK, Attwood J, Burger M, et al. DNA methylation profiling of human chromosomes 6, 20 and 22. Nat Genet. 2006;38:1378–1385.

8. Sati S, Tanwar VS, Kumar KA, Patowary A, Jain V, Ghosh S, et al. High resolution methylome map of rat indicates role of intragenic DNA methylation in identification of coding region. PLoS One. 2012;7(2):e31621.

9. Sandra Cortijo René, Wardenaar, Maria Colomé-Tatché, Frank Johannes, Vincent Colot Sandra Cortijo, et al. Genome-Wide Analysis of DNA Methylation in Arabidopsis Using MeDIP-Chip. Plant Epigenetics and Epigenomics.2014;1112:125–149.

10. Buckingham M, Vincent SD. Distinct and dynamic myogenic populations in the vertebrate embryo. Curr Opin Genet Dev. 2009;19(5):444–53.

11. Buckingham M. Skeletal muscle formation in vertebrates. Curr Opin Genet Dev. 2001;11:440–8.

12. Larsen F, Gundersen G, Lopez R, Prydz H: Cpg Islands as gene markers in the human genome. Genomics. 1992; 13(4): 1095–1107.

13. Plass C, Soloway PD: DNA methylation, imprinting and cancer. Eur J Hum Genet. 2002; 10(1):6–16.

14. Su ZX, Xia JF, Zhao ZM. Functional complementation between transcriptional methylation regulation and post-transcriptional microRNA regulation in the human genome. BMC Genomics. 2011; 12(Suppl 5):S15.

15. Frommer M, McDonald LE, Millar DS, Collis CM, Watt F, et al. A genomic sequencing protocol that yields a positive display of 5-methylcytosine residues in individual DNA strands. Proc Natl Acad Sci U S A. 1992; 89:1827–1831.

16. Wang S, Lv J, Zhang L, Dou J. MethylRAD: a simple and scalable method for genome-wide DNA methylation profiling using methylation dependent restriction enzymes. Open Biol. 2015; 150130:5.

17. Cohen-Karni D, Xu D, Apone L,Fomenkov A,Sun Z, Davis PJ, et al. The MspJI family of modification-dependent restriction endonucleases for epigenetic studies. Proc Natl Acad Sci U S A. 2011;108,11040–11045.

18. Zhang Y, Liu J, Fu W, Xu W, Zhang H, Chen S, et al. Comparative Transcriptome and DNA methylation analyses of the molecular mechanisms underlying skin color variations in Crucian carp (Carassius carassius L.). BMC Genet. 2017;18(1):95

19. Li H, Yuan J, Wu M, Han Z, Li L, Jiang H, et al. Transcriptome and DNA methylome reveal insights into yield heterosis in the curds of broccoli (Brassica oleracea L var. italic) BMC Plant Biol. 2018;18(11):168.

20. Krueger F, Andrews SR. Bismark: a flexible aligner and methylation caller for bisulfite-Seq applications. Bioinformatics. 2011;27(1): 1571–2.

21. Weber M, Davies JJ, Wittig D, Oakeley EJ, Haase M, Lam WL, Schuebeler D: Chromosome-wide and promoter-specific analyses identify sites of differential DNA methylation in normal and transformed human cells. Nat Genet. 2005;37(8):853–862.

22. Li Q, Li N, Hu X, Li J, Du Z, Chen L, Yin G, Duan J, Zhang H, Zhao Y: Genome-wide mapping of DNA methylation in chicken. Plos One. 2011;6(5):e19428.

23. Zhang X, Yazaki J, Sundaresan A, Cokus S, Chan SW-L, Chen H, Henderson IR, Shinn P, Pellegrini M, Jacobsen SE: Genome-wide high-resolution mapping and functional analysis of DNA methylation in arabidopsis. Cell. 2006;126(6):1189–1201.

24. Ball MP, Li JB, Gao Y, Lee JH, LeProust EM, Park I-H, et al. Targeted and genome-scale strategies reveal gene-body methylation signatures in human cells. Nat Biotechnol. 2009, 27(4):361–368.

25. Laurent L, Wong E, Li G, Huynh T, Tsirigos A, Ong CT, et al. Dynamic changes in the human methylome during differentiation. Genome Res. 2010;20(3):320–31.

26. Feng S, Cokus ST, Zhang X, Chen PY, Bostick M, Goll MG, et al. Conservation and divergence of methylation patterning in plants and animals. Proc Natl Acad Sci U S A. 2010;107(19):8689–94.

27. Long J, Jing Z, Xia YD, Lou P, Chen L, Wang HM, et al. Genome-wide DNA methylation changes in skeletal muscle between young and middle-aged pigs. BMC Genomics. 2014;15(1):653.

28. Shukla S, Kavak E, Gregory M, Imashimizu M, Shutinoski B, Kashlev M, et al. CTCF-promoted RNA polymerase II pausing links DNA methylation to splicing. Nature. 2011;479(7371):74–79.

29. Lorincz MC, Dickerson DR, Schmitt M, Groudine M. Intragenic DNA methylation alters chromatin structure and elongation eficiency in mammalian cells. Nat. Struct. Mol. Biol. 2004;11, 1068–1075.

30. Klose RJ, Bird AP. Genomic DNA methylation: the mark and its mediators. Trends Biochem. Sci. 2006;31,89–97.

31. Suzuki MM, Bird A. DNA methylation landscapes: provocative insights from epigenomics. Nat Rev Genet. 2008;9(6):465–76.

32. Li MZ, Wu HL, Luo ZG, Xia YD, Guan JQ, Wang T, et al: An atlas of DNA methylomes in porcine adipose and muscle tissues. Nature communications. 2012;3:850.

33. Klose RJ, Bird AP: Genomic DNA methylation: the mark and its mediators. Trends Biochem Sci. 2006;31(2):89–97

34. Lorincz MC, Dickerson DR, Schmitt M, Groudine M. Intragenic DNA methylation alters chromatin structure and elongation efficiency in mammalian cells. Nat Struct Mol Biol. 2004;11(11):1068–75

35. Rountree MR, Selker EU. DNA methylation inhibits elongation but not initiation of transcription in Neurospora crassa. Genes Dev. 1997;11(18):2383–95.

36. Zhu H, Wang G, Oian J. Transcription factors as readers and effectors of DNA methylation Nat Rev Genet. 2016;17(9):551–65.

37. Harel NY, Strittmatter SM. Can regenerating axons recapitulate developmental guidance during recovery from spinal cord injury? Nat Rev Neurosci. 2006;7(8):603–16.

38. Zykovich A, Hubbard A, Flynn JM, Tarnopolsky M, Fraga MF, Kerksick C, et al. Genome-wide DNA methylation changes with age in disease free human skeletal muscle. Aging Cell. 2014;13(21):360–6.

39. Zihni C, Mills C, Matter K, Balda MS. Tight junctions: from simple barriers to multifunctional molecular gates. Nat Rev Mol Cell Biol. 2016;17(9):564–80.

40. Potter E, Bergwitz C, Brabant G. The cadherin-catenin system: implications for growth and differentiation of endocrine tissues. Endocr Rev. 1999;20(2):207–39.

41. Bazzoni G, Dejana E. Endothelial cell-to-cell junctions: molecular organization and role in vascular homeostasis. Physiol Rev. 2004;84(3):869–901.

42. Tran TH, Shi X, Zaia J, Ai X. Heparan Sulfate 6-O-endosulfatases (Sulfs) Coordinate the Wnt Signaling Pathways to Regulate Myoblast Fusion during Skeletal Muscle Regeneration. J Biol Chem. 2012;287(39):32651–32664.

43. Cisternas P, Henriquez JP, Brandan E, Inestrosa NC. Wnt Signaling in Skeletal Muscle Dynamics: Myogenesis, Neuromuscular Synapse and Fibrosis. Mol Neurobiol. 2014;49(1):574–89.

44. van Amerongen R, Berns A. Knockout mouse models to study Wnt signal transduction. Trends Genet. 2006; 22(12):678–689.

45. Roffers-Agarwal, J, Xanthos, JB, Miller JR. Regulation of actin cytoskeleton architecture by Eps8 and Abi1. BMC Cell Biol. 2005;6:36.

46. Tapscott SJ. The circuitry of a master switch: Myod and the regulation of skeletal muscle gene transcription. Development, 2005,132(12):2685–2695.

47. Montarras D1, Chelly J, Bober E, Arnold H, Ott MO, Gros F, et al. Developmental patterns in the expression of Myf5, MyoD, myogenin, and MRF4 during myogenesis. New Biol. 1991;3(6):592–600.

48. Laclef C, Hamard G, Demignon J, Souil E, Houbron C, Maire P. Altered myogenesis in Six1-deficient mice. Development. 2003;130(10):2239–52.

49. Grifone R, Demignon J, Houbron C, Souil E, Niro C, Seller MJ, et al. Six1 and Six4 homeoproteins are required for Pax3 and Mrf expression during myogenesis in the mouse embryo. Development. 2005;132(9):2235–49.

50. Richard AF, Demignon J, Sakakibara I, Pujol J, Favier M, Strochlic L, et al. Genesis of muscle fiber-type diversity during mouse embryogenesis relies on Six1 and Six4 gene expression. Dev Biol. 2011;359(2):303–20.

51. Liu Y, Chakroun I, Yang D, Horner E, Liang J, Aziz A, et al. Six1 Regulates MyoD Expression in Adult Muscle Progenitor Cells. PLoS One. 2013;8(6):e67762.

52. Gianakopoulos PJ, Mehta V, Voronova A, Cao Y, Yao Z, Coutu J, et al. MyoD directly up-regulates premyogenic mesoderm factors during induction of skeletal myogenesis in stem cells. J Biol Chem. 2011;286(4):2517–25.

53. Karczewska-Kupczewska M, Stefanowicz M, Matulewicz N, Nikolajuk A, Straczkowski M. Wnt Signaling Genes in Adipose Tissue and Skeletal Muscle of Humans With Different Degrees of Insulin Sensitivity. J Clin Endocrinol Metab. 2016 Aug; 101(8):3079–87. doi: 10.1210/jc.2016-1594.

54. Wang S, Liu P, Lv J, Li Y, Cheng T, Zhang L, et al. Serial sequencing of isolength RAD tags for cost-efficient genome-wide profiling of genetic and epigenetic variations. Nature Protocols, 2016, 11 (11):2189–2200.

55. Li R, Yu C, Li Y, Lam TW, Yiu SM, Kristiansen K. SOAP2: an improved ultrafast tool for short read alignment. Bioinformatics. 2009;25(15):1966–7.

56. Robinson MD1, McCarthy DJ, Smyth GK. edgeR: a Bioconductor package for differential expression analysis of digital gene expression data. Bioinformatics. 2010;26 (1): 139–40.

57. Cohen-Karni D, Xu D, Apone L, Fomenkov A, Sun Z, Davis PJ, et al. The MspJI family of modification-dependent restriction endonucleases for epigenetic studies. Proc Natl Acad Sci U S A 2011;108(27):11040–5.

58. Patel RK, Jain M. NGS QC Toolkit: A toolkit for quality control of next generation sequencing data. PLoS ONE. 2012; 7 (2): e30619.

59. Trapnell C, Williams BA, Pertea G, Mortazavi A, Kwan G, van Baren MJ, et al. Transcript assembly and quantification by RNA-Seq reveals unannotated transcripts and isoform switching during cell differentiation. Nat Biotechnol. 2010;28(5):511–5.

60. Langmend B, Salzberg SL. Fast gapped-read alignment with Bowtie 2. Nat Methods. 2012;9(4):357–9.

61. Mortazavi A, Williams BA, McCue K, Schaeffer L, Wold B. Mapping and quantifying mammalian transcriptomes by RNA-Seq. Nat Methods. 2008;5(7):621–8.

62. Kanehisa M, Araki M, Goto S, Hattori M, Hirakawa M, Itoh M, et al. KEGG for linking genomes to life and the environment. Nucleic Acids research. 2008;36:480–4.

63. Livak KJ, Schmittgen TD: Analysis of relative gene expression data using real-time quantitative PCR and the 2(-Delta Delta C(T)) Method. Methods. 2001;25(4):402–8.

